# Radiochemical synthesis and evaluation in nonhuman primates of 3-[^11^C]methoxy-4-aminopyridine: a novel PET tracer for imaging potassium channels in the CNS

**DOI:** 10.1101/2020.12.11.421578

**Authors:** Nicolas J. Guehl, Ramesh Neelamegam, Yu-Peng Zhou, Sung-Hyun Moon, Maeva Dhaynaut, Georges El Fakhri, Marc D. Normandin, Pedro Brugarolas

## Abstract

Demyelination, the loss of the protecting sheath of neurons, contributes to disability in many neurological diseases. In order to fully understand its role in different diseases and to monitor treatments aiming at reversing this process, it would be valuable to have PET radiotracers that can detect and quantify molecular changes involved in demyelination such as the uncovering and upregulation of the axonal potassium channels K_v_1.1 and K_v_1.2. Carbon-11 labeled radiotracers present the advantage of allowing for multiple scans on the same subject in the same day. Here, we describe [^11^C|3MeO4AP, a novel ^11^C-labeled version of the K^+^ channel tracer [^18^F]3F4AP, and characterize its imaging properties in two nonhuman primates including a monkey with a focal brain injury sustained during a surgical procedure three years prior to imaging. Our findings show that [^11^C]3MeO4AP is brain permeable, metabolically stable and has high plasma availability. When compared with [^18^F]3F4AP, [^11^C]3MeO4AP shows very high correlation in volumes of distribution (*V_T_*) confirming a common target. [^11^C]3MeO4AP shows slower washout than [^18^F]3F4AP suggesting stronger binding. Finally, similar to [^18^F]3F4AP, [^11^C]3MeO4AP is highly sensitive to the focal brain injury. All these features make it a promising radioligand for imaging demyelinated lesions.

## Introduction

Demyelination – the loss of the insulating sheath that surrounds axons – is a major contributor to disability in multiple sclerosis (MS)^1^ and likely in other diseases such as stroke^2^, traumatic brain and spinal cord injuries^3,4^ and Alzheimer’s disease^5^. In order to understand the role of demyelination in different neurological diseases and to be able to monitor therapies aiming to promote remyelination, it is critical to develop noninvasive imaging tools that are sensitive to demyelination^6,7^.

To that end, we previously described [^18^F]3F4AP, a radiofluorinated analog of the multiple sclerosis drug 4AP, as a promising radioligand for imaging demyelination^8–10^. [^18^F]3F4AP binds to axonal voltage-gated K^+^ (K_v_) channels, which increase in expression and become more accessible after demyelination^11–15^. Given the significant changes in K^+^ channels with demyelination, we previously hypothesized that [^18^F]3F4AP could be used to detect demyelinated lesions using positron emission tomography (PET) and, subsequently, confirmed this experimentally in rodent models of demyelination^8^. More recently, we evaluated [^18^F]3F4AP in nonhuman primates (NHPs) and demonstrated that it has favorable imaging properties and is sensitive to a focal brain injury^16^.

While ^18^F-labeled tracers (*t*_1/2_ = 110 min) can be produced remotely and distributed regionally, there are instances where having a shorter-lived radiotracer can be advantageous. For example, ^11^C-labeled tracers (*t*_1/2_ = 20.4 min) allow performing two scans on the same subject in the same day, which facilitates the assessment of test/retest variability, conducting paired baseline-blocking studies, comparing two radiotracers, or acquiring data with two complementary radiotracers. Furthermore, developing different ^18^F- and ^11^C-labeled analogs of a given compound provides the opportunity to perform structure-activity relationship studies, which can potentially result in the identification of radioligands with improved physicochemical properties. Thus, in a recent work, we investigated four novel derivatives of 4AP as potential candidates for PET imaging of demyelination^17^. Among those, 3-methoxy-4-aminopyridine (3MeO4AP) was found to bind to Shaker K_v_ channel with similar affinity as 4AP and 3F4AP and demonstrated favorable physicochemical properties. Its p*K*_a_ value was found to be 9.18 indicating that at physiological pH this compound is mostly in its protonated form, the required form for binding to the channel. In terms of lipophilicity, 3MeO4AP was found to have an octanol/water partition coefficient at pH 7.4 (logD) of −0.76 suggesting that it may penetrate the blood-brain barrier (BBB) faster than 4AP (logD = −1.48) but slower than 3F4AP (logD = 0.41). Taken together, these *in vitro* results suggest that 3MeO4AP is a promising candidate for PET imaging.

In the present work, we present the novel ^11^C-labeled radiotracer 3-[^11^C]methoxy-4-aminopyridine ([^11^C]3MeO4AP), and its evaluation in NHPs. Specifically, we describe its radiochemical synthesis via [^11^C]methylation, its characterization in NHPs including pharmacokinetic modeling with arterial input functions, and the direct comparison with [^18^F]3F4AP using a graphical method previously reported by Guo et al^18^.

## Results and Discussion

### 3-[^11^C]methoxy-4-aminopyridine ([^11^C]3MeO4AP) synthesis by direct ^11^C-methylation

Radiochemical synthesis of [^11^C]3MeO4AP (**2**) was accomplished via [^11^C]methylation of 3-hydroxy-4-aminopyridine (**1**) with [^11^C]methyl iodide in DMSO at 80 °C under basic conditions (**Figure 1A**). A summary of the conditions tested are shown in **Sup. Table 1**. Using this method, [^11^C]3MeO4AP (**2**) was produced with high molar activity (1.4 ± 0.1 Ci/μmol), high radiochemical purity (> 98.5%) and 15 ± 4% (*n* = 6) non-decay corrected radiochemical yield (**Figure 1B** and **C**). The total time from end of bombardment to reformulation was *ca*. 40 min.

**Figure 1.**
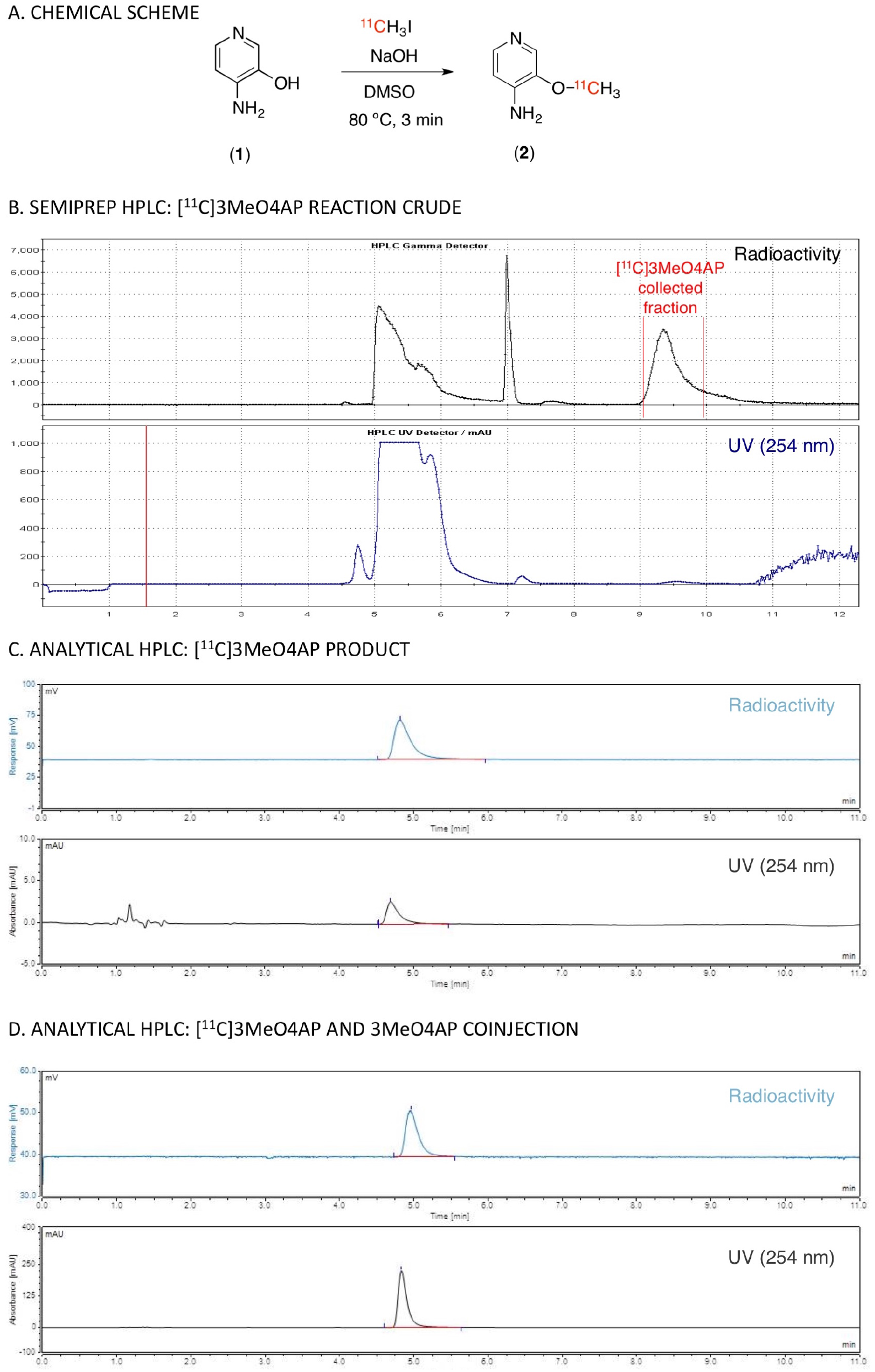
Radiochemical synthesis of [^11^C]3MeO4AP. **(A)** Chemical reaction scheme. (**B**) Semipreparative HPLC (high performance liquid chromatography) purification (radio and UV chromatograms). (**C**) Analytical HPLC (radio and UV chromatograms) of tracer only. (**D**) Analytical HPLC (radio and UV chromatograms) of tracer plus reference standard.

### [^11^C]3MeO4AP in nonhuman primates: Analysis of arterial blood

Immediately after producing [^11^C]3MeO4AP, the radiotracer was administered intravenously as a 3 min slow bolus to nonhuman primates (NHPs) positioned inside a PET/CT scanner. During the scan, serial arterial samples were collected and analyzed for radioactivity concentration in whole-blood (WB) and plasma (PL), radiometabolites and plasma free fraction (*f_p_*). Total radioactivity in WB and PL peaked quickly and cleared rapidly after the 3 min venous infusion of [^11^C]3MeO4AP (**Figure 2A** and **B**). The ratio of WB to PL concentration reached equilibrium within 1 min and was close to unity (WB:PL at 1-min post-injection across all studies = 1.21 ± 0.04). RadioHPLC metabolite measurements showed minimal metabolic degradation with 86.2 ± 4.6% parent remaining after 60 min post injection (**Figure 2C** and **Supplemental Figure 1**). Finally, *f_p_* measurements showed minimal binding to plasma proteins (*f_p_* = 100% in Monkey 1 and 96% in Monkey 2, measured in triplicate for n = 1 imaging experiment per monkey).

**Figure 2:**
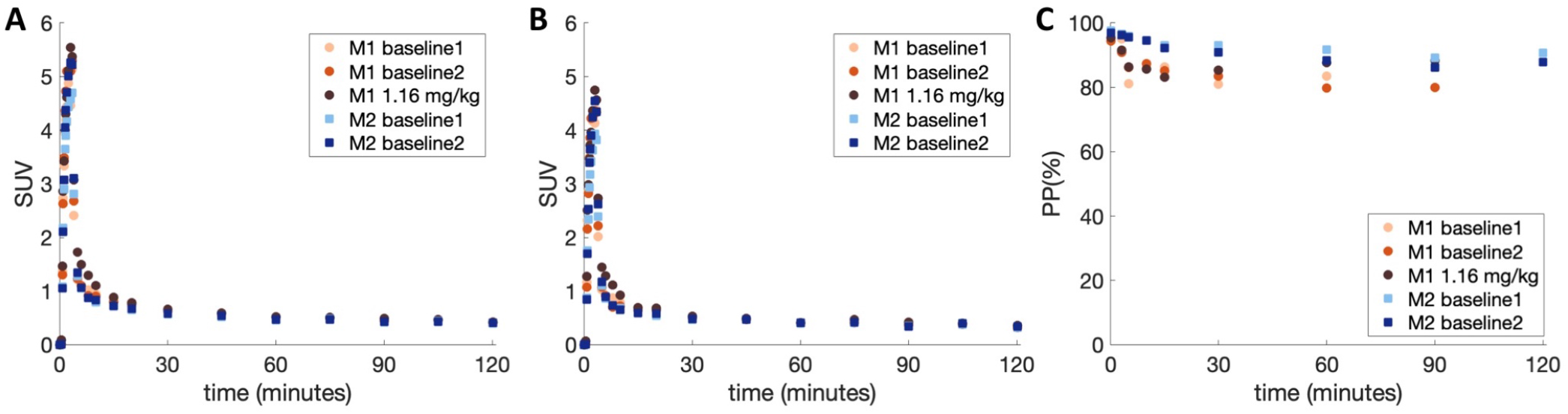
[^11^C]3MeO4AP in arterial blood. (**A**) Time course in whole blood (WB). (**B**) Time course in plasma (PL). (**C**) [^11^C]3MeO4AP percent parent in plasma (%PP). In the legends, M1 and M2 refer to Monkey 1 and 2, respectively.

### [^11^C]3MeO4AP in nonhuman primates: Brain uptake and pharmacokinetic modeling

[^11^C]3MeO4AP entered the brain readily and peaked relatively quickly. 20 min post-injection the standardized uptake value (SUV) in gray matter areas was greater than 2 and around 1.5 in white matter. Uptake and kinetics were relatively heterogeneous across brain regions. **Figure 3A** shows the time activity curves (TACs) from baseline studies acquired in each monkey and **Figure 3B** the SUV images calculated from 45 to 90 min post tracer injection (SUV_45-90min_).

**Figure 3:**
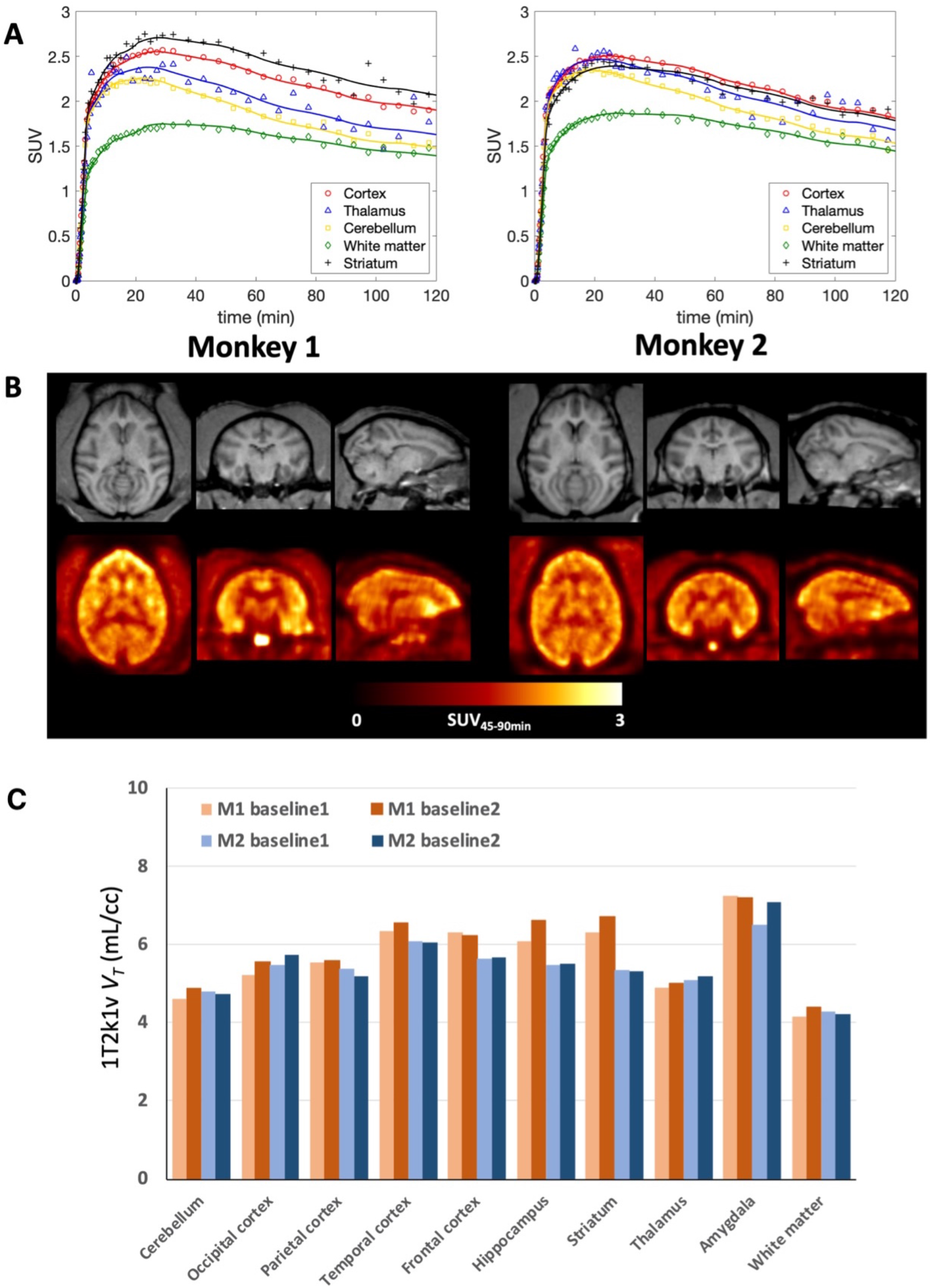
[^11^C]3MeO4AP in the primate brain. (**A**) Brain time activity curves and model fits using a reversible 1T model with the vascular contribution *v* included as a model parameter (1T2k1v). (**B**) Individual MRI MEMPRAGE (top) and PET SUV images calculated from 45 to 90 min post tracer injection (SUV_45-90min_) (bottom). Images are presented in the MRI NIMH Macaque template (NMT) space^21^. For each panel (A and B), left corresponds to data acquired in Monkey 1 and right corresponds to Monkey 2. For each monkey, data shown correspond to a single baseline study. (**C**) Regional *V_T_* values for each baseline scan and animal.

Visually, brain TACs were well described by both reversible one tissue (1T) and two tissue (2T) compartment models for the entire 120 min of scan acquisition. However, in this work 90 min of data was used as the reference scan duration for the kinetic analyses because of increased statistical noise in late PET images as well as in the parent fraction measurements. According to the Akaike information criterion (AIC)^19^, over all TACs and studies the preferred model was a reversible 1T model with the vascular contribution *v* included as a model parameter (1T2k1v) (AIC_weight,median_ = 0.339), followed by a reversible 2T model with the vascular contribution *v* included as a model parameter (2T4k1v) (AIC_weight,median_ = 0.128). *V_T_* values estimated from the 1T2k1v and 2T4k1v models were stable and in very good agreement (*V_T,2T4k1v_* = 0.99 x *V_T,1T2k1v_* + 0.08; *r* = 0.99, *p* < 0.0001; mean difference = 0.03 ± 0.1 mL/cc, average measure intraclass correlation coefficient ICC = 0.996 with CI95% of [0.993, 0.998]). Since the 1T2k1v model adequately described the kinetics of [^11^C]3MeO4AP (**Figure 3A**) and the more complex 2T4k1v model was not statistically justified, we chose the 1T2k1v model to quantify [^11^C]3MeO4AP signal. The quantitative analysis confirmed the good brain penetration of the radiotracer with an average *K_1_* value of 0.16 ± 0.01 mL/min/cc for the whole brain, as measured across all studies. **Figure 3C** shows bar plots of the average *V_T_* values of baseline scans for each animal. The lowest *V_T_* value was measured in the white matter and *V_T_* measurements were higher in cortical areas. The highest *V_T_* was found in the amygdala. Test-retest (TRT) was excellent in Monkey 2 (mean TRT = 1.0 ± 3.6% across brain regions, average measure ICC = 0.976 with CI95% of [0.908, 0.994]) and so was the reproducibility of *V_T_* measurements for the two baseline scans acquired in Monkey 1 three months apart (mean difference = 3.9 ± 3.3% across brain regions, average measure ICC = 0.978 with CI95% of [0.722, 0.996]). Across all baseline studies and brain regions, the average coefficient of variation (COV) was 5.6 ± 3.3%, with the lowest COV measured in the white matter (2.4%) and highest COV measured in the striatum (11.7%). **Table 1** shows *V_T_* values obtained for all studies. Coadministration of [^11^C]3MeO4AP and nonradioactive 3MeO4AP (1.16 mg/kg) did not show a reduction in *V_T_*, likely because the dose was below the pharmacological dose. These findings are consistent with previous studies with [^18^F]3F4AP, which showed that administration of similar doses of 3F4AP did not cause a reduction in *V_T_* likely due to a compensatory mechanism by which when some channels become occupied others transition to the open and bindable conformation^16^. We did not attempt coadministration of higher doses of nonradioactive 3MeO4AP as previous studies have shown that high doses of 4AP and related compounds can cause seizures^8,20^.

**Table 1:** Regional 1T2k1v *V_T_* estimates obtained from all studies

| ROI | Monkey 1 |  |  | Monkey 2 |  |
| --- | --- | --- | --- | --- | --- |
|  | Baseline 1 | Baseline 2 | 1.16 mg/kg | Baseline 1 | Baseline 2 |
| Cerebellum | 4.59 | 4.89 | 4.55 | 4.79 | 4.74 |
| Occipital Cortex | 5.21 | 5.55 | 5.21 | 5.47 | 5.74 |
| Parietal Cortex | 5.53 | 5.59 | 5.40 | 5.37 | 5.18 |
| Temporal Cortex | 6.35 | 6.54 | 6.47 | 6.09 | 6.06 |
| Frontal Cortex | 6.30 | 6.23 | 6.40 | 5.63 | 5.66 |
| Hippocampus | 6.07 | 6.60 | 6.13 | 5.47 | 5.51 |
| Striatum | 6.30 | 6.71 | 6.51 | 5.36 | 5.32 |
| Thalamus | 4.90 | 5.02 | 4.99 | 5.09 | 5.19 |
| Amygdala | 7.23 | 7.20 | 7.00 | 6.50 | 7.08 |
| White matter | 4.17 | 4.40 | 4.21 | 4.27 | 4.21 |
| Whole brain | 5.16 | 5.40 | 5.19 | 4.99 | 5.02 |

*V_T_* estimates were stable when truncating the data down to 60 min. *V_T_* values obtained using only 60 min of data were in excellent agreement with those obtained using 90 min of data (mean difference = 0.007 ± 0.224 mL/cc, average measure ICC = 0.984 with CI95% of [0.972,0.991]). Likewise, and despite the increased statistical noise in the measurements late in the scan, *V_T_* estimates obtained when using 120 min of data were in excellent agreement with those obtained using 90 min of data (mean difference = 0.000 ± 0.125 mL/cc, average measure ICC = 0.994 with CI95% of [0.989,0.997]).

Logan plots linearized very well by a *t*\* of 10 min and resulted in estimates of *V_T_* that were highly correlated with those obtained from the 1T2k1v model despite some small underestimation (*V_T,Logan_* = 0.91 x *V_T,1T2k1v_* + 0.27; *r* = 0.99, *p* < 0.0001; mean difference = −0.22 ± 0.10 mL/cc, average measure ICC = 0.977 with CI95% of [0.149,0.995]). **Figure 4** shows Logan plots as well as parametric maps of *V_T_* obtained for each animal. Similar to the observation made for the Logan graphical method, using a *t*\* of 10 min the multilinear analysis 1 (MA1) regression model demonstrated good agreement with the full compartmental analysis with a small underestimation (*V_T,MA1_*= 0.96 x *V_T,1T2k1v_* + 0.06; *r* = 0.99, *p*<0.0001; mean difference = −0.19 ± 0.09 mL/cc, average measure ICC = 0.984 with CI95% of [0.223,0.996]).

**Figure 4:**
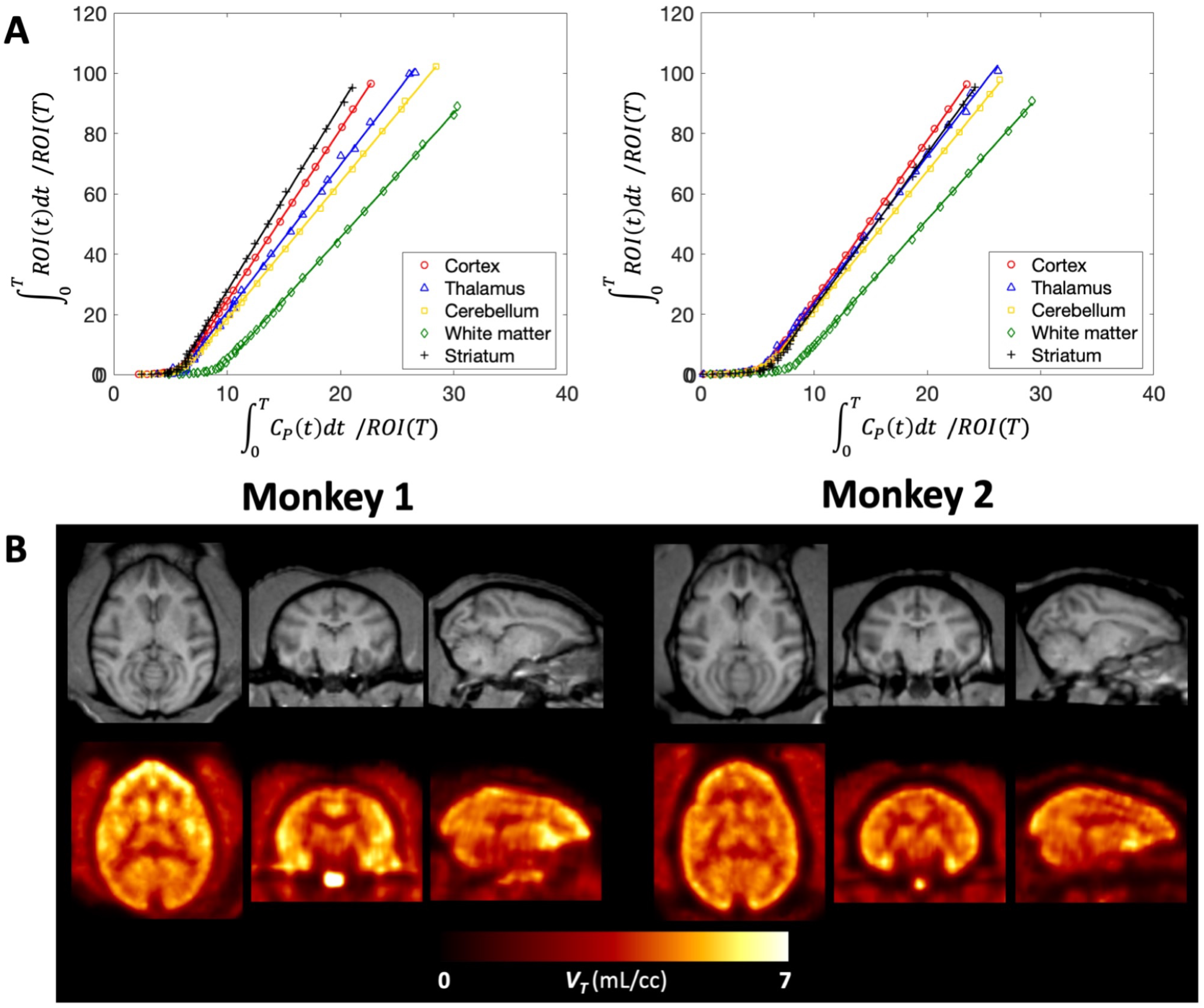
Graphical analysis of [^11^C]3MeO4AP in the primate brain. (**A**) Logan plots using a 10-min *t*\*. (**B**) Individual MRI MEMPRAGE images (top) and PET parametric images of total volume of distribution *V_T_* (bottom), after registration into the NMT space. For each panel (A and B), left corresponds to data acquired in Monkey 1 and right corresponds to Monkey 2. For each monkey, data shown correspond to a single baseline study. Logan plots (A) and Logan *V_T_* parametric maps were generated using 90 minutes of data.

### [^11^C]3MeO4AP in nonhuman primates: Sensitivity to a focal traumatic brain injury

One of the monkeys we imaged had sustained a minor intracranial injury during a craniotomy procedure three years prior to imaging. The location of the craniotomy site can be clearly seen on the CT images (**Figure 5A top**). PET images of the region of the brain right under the burr hole on the skull showed a marked increase in [^11^C]3MeO4AP signal (**Figure 5A bottom**). Quantitative analysis showed a 60.3% increase in *V_T_* in this region compared to the contralateral side (*V_T_* lesion = 7.29 mL/cc, *V_T_* contra = 4.55 mL/cc; TRT = 0.9%, n = 2) and slower kinetics (**Figure 5B**). This change could not be attributed to increase in perfusion as the 1T2k1v model actually showed a decrease in *K_1_* (−38.7% on average between the two baseline scans) in this region compared to the contralateral side. Despite the slower kinetics in the lesion, *V_T_* estimates were relatively stable when truncating the acquisition down to 60 min with differences in *V_T_* within 10% when compared to 90 min of data.

**Figure 5:**
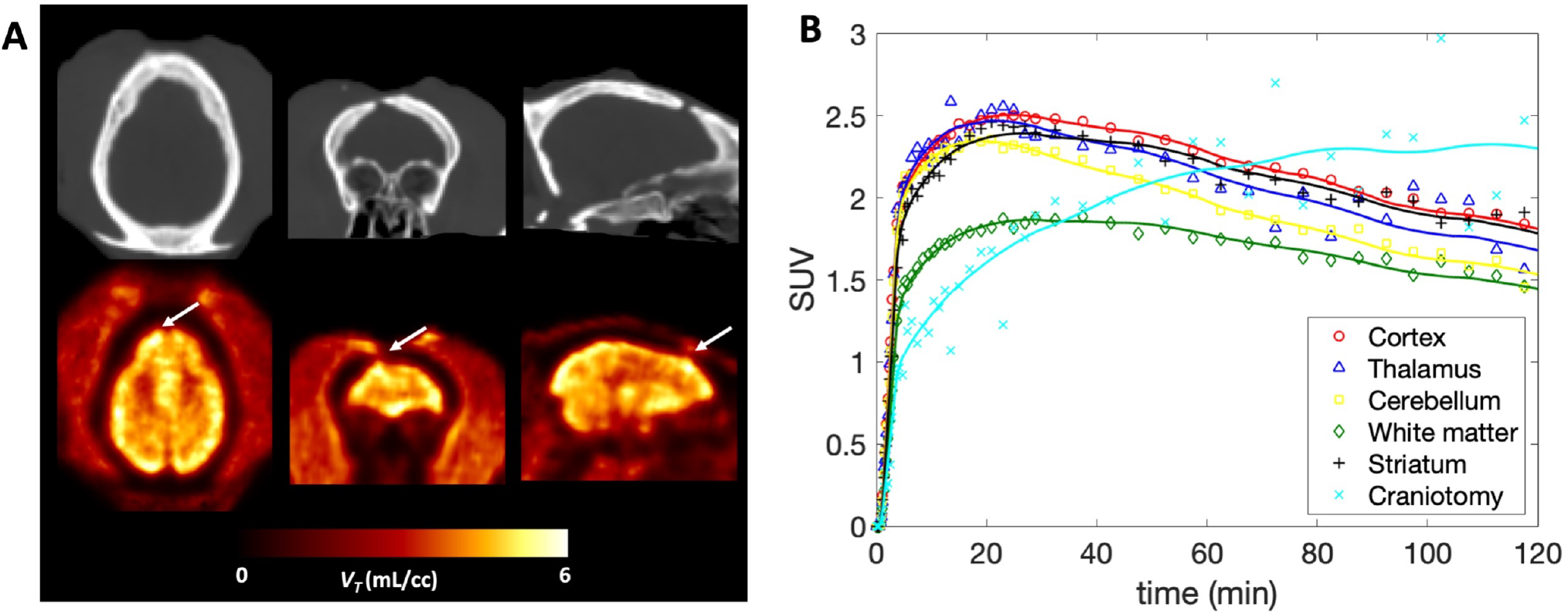
[^11^C]3MeO4AP in the focal brain injury. (**A**) CT and [^11^C]3MeO4AP parametric images of *V_T_* with axial, coronal and sagittal view showing the injury. Arrows indicate the location of the lesion. (**B**) Time activity curves including the lesion in light blue of Monkey 2 showing distinct kinetics compared to other regions.

### Direct comparison between [^11^C]3MeO4AP and [^18^F]3F4AP using a graphical method

We recently reported the pharmacokinetic evaluation of [^18^F]3F4AP in the same two animals^16^. Compared to [^18^F]3F4AP, [^11^C]3MeO4AP displayed more heterogeneity across brain regions and slower kinetics. The rank order of *V_T_* values across brain regions was largely consistent between the two tracers. Direct comparison of the regional *V_T_* values of [^11^C]3MeO4AP and [^18^F]3F4AP in both animals showed a strong linear relationship (**Figure 6A**: *r* = 0.97, *p*<0.0001 for Monkey 1; **Figure 6B**: *r* = 0.85, *p*=0.0017 for Monkey 2; **Figure 6C**: *r* = 0.93, *p*<0.0001 for Monkey 1 and 2 combined on the same plot) indicating that the two ligands bind to a common target. The linear regressions showed negative *y*-intercepts (**Figure 6A**: y = −1.63, *p*=0.0312 for Monkey 1; **Figure 6B**: y = −0.58, *p*=0.6693 for Monkey 2; **Figure 6C**: y = −1.27, *p*=0.0561 for both animals combined) suggesting that [^11^C]3MeO4AP has higher specific binding than [^18^F]3F4AP 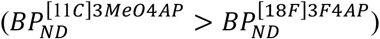. After swapping the axes of plots in **Figure 6**, *y*-intercept of all plots became positive (y = 0.62, *p* = 0.0031 for Monkey 1; y = 0.78, *p*=0.0444 for Monkey 2; y = 0.67, *p*=0.0003 for both animals combined, graphs not shown) thus preserving the rank order of 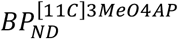 *and* 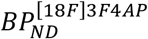. This result indicates that the determination of the rank order of *BP_ND_* between [^11^C]3MeO4AP and [^18^F]3F4AP from the y-intercept is not affected to a significant level by noise on the axis^18^, as standard linear regression assumes that data plotted on the x-axis is noiseless. Lastly, we estimated the *in vivo K_D_* ratio between [^11^C]3MeO4AP and [^18^F]3F4AP using the *f_p_* measurements of both tracers. The *in vivo* affinity ratios 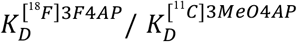 were 2.82 for Monkey 1 (**Figure 6A**), 2.58 for Monkey 2 (**Figure 6B**) and 2.78 when combining Monkey 1 and 2 on the same graph (**Figure 6C**), thus suggesting that [^11^C]3MeO4AP has higher *in vivo* affinity than [^18^F]3F4AP for K_v_ channels.

**Figure 6:**
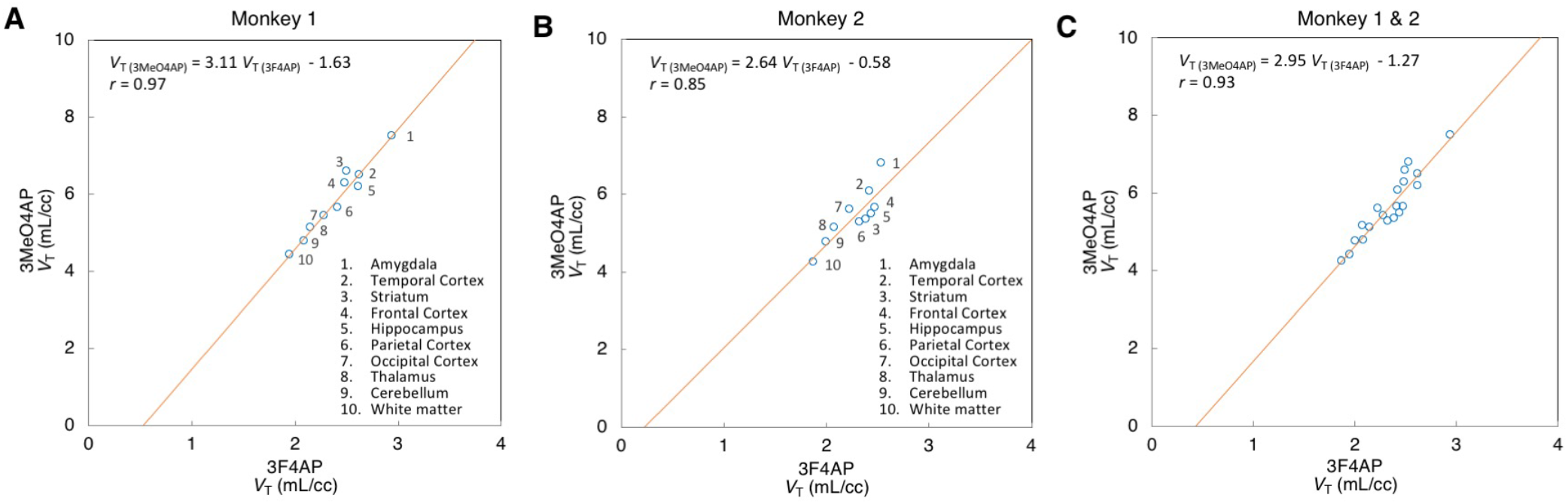
Scatter plots comparing the *V_T_* values obtained for [^11^C]MeO4AP and [^18^F]3F4AP in Monkey 1 (A), Monkey 2 (B) and both together (C). For each radiotracer the plotted *V_T_* values represent the average of the two baseline scans. Paired comparisons are plotted for the regional *V_T_* estimates in occipital cortex, parietal cortex, temporal cortex, frontal cortex, hippocampus, amygdala, striatum, thalamus, white matter and whole cerebellum. Red lines represent the linear regression lines.

### Summary and discussion

Demyelination is emerging as an important pathological feature in many neurological diseases in addition to multiple sclerosis. These include traumatic brain and spinal cord injuries, stroke, dementias and psychiatric disorders. Given the lipophilic nature of the myelin sheath, demyelinated lesions are usually detectable on MRI. However, there are some limitations for imaging demyelination using MR sequences, including a lack of specificity^22^, the difficulty of detecting grey matter lesions where myelin is discontinuous and sparse^23^ and the challenge of obtaining quantitative measurements^24^. Given these limitations, the use of PET radioligands that target proteins differentially expressed in demyelinated lesions appears to be a highly promising approach^6^. To that end, we previously developed [^18^F]3F4AP (a radiofluorinated analog of the multiple sclerosis drug 4AP) as an ^18^F-labeled PET ligand for voltage-gated potassium channels (K_v_1.1 and K_v_1.2)^8^. K_v_1.1 and K_v_1.2 are a relevant target as these channels significantly increase in expression and in accessibility upon demyelination^12,13,25^. In our previous work, we showed that [^18^F]3F4AP can detect demyelinated lesions in rodent models of MS^8^ and, more recently, that it has favorable pharmacokinetic properties in non-human primates and is highly sensitive to a focal brain injury^16^. In a parallel but related effort, we also investigated four novel derivatives of 4AP as potential candidates for imaging and/or therapy, among which 3MeO4AP was found to possess good binding affinity as well as suitable basicity (essential for binding) and lipophilicity (essential for brain penetration)^17^.

In the present work, we described the radiochemical synthesis of [^11^C]3MeO4AP via standard ^11^C-methylation using [^11^C]methyl iodide (**Figure 1**). Subsequent imaging studies in NHPs showed suitable pharmacokinetics both in plasma and in brain. Analyses of arterial plasma samples demonstrated a high metabolic stability, minimal plasma protein binding, and relatively rapid plasma clearance (**Figure 2**). [^11^C]3MeO4AP readily entered the monkey brain and showed relatively heterogenous uptake and kinetics across brain regions (**Figure 3**). Highest uptake was seen in the amygdala and lowest in the white matter. A 1-tissue compartment model with the vascular contribution *v* included as a model parameter (1T2k1v) was found to be sufficient to describe the brain time activity curves in all brain regions and provided robust estimates of the total volume of distribution (*V_T_*) (**Figure 4**). Test-retest performed on the same day in Monkey 2 was excellent and so was the reproducibility of *V_T_* estimates for the two baseline scans acquired in Monkey 1, thus demonstrating the basic reproducibility of [^11^C]3MeO4AP quantitative measurements. Simplified methods such as the Logan and MA1 graphical methods provided *V_T_* measurements that were in good agreement with those obtained from the full compartmental analysis. Similar to [^18^F]3F4AP, [^11^C]3MeO4AP was found to be highly sensitive to a focal traumatic injury that occurred 3 years prior to imaging (**Figure 5**).

When compared to [^18^F]3F4AP, [^11^C]3MeO4AP demonstrated slower brain penetration, which can be quantitatively assessed from the *K_1_* estimates (*K*_*1*[^11^C]3MeO4AP_ = 0.16 ± 0.01 mL/min/cc *vs*. *K*_*1*[^18^F]3F4AP_ = 0.45 ± 0.10 mL/min/cc, measured in whole brain across baseline studies), followed by slower washout. The differences in brain penetration can be attributed to the lower lipophilicity of [^11^C]3MeO4AP compared to [^18^F]3F4AP (logD at pH 7.4 = −0.76 vs. 0.41). The slower washout is consistent with stronger binding of [^11^C]3MeO4AP to K_v_ channels in the CNS. This was quantitatively assessed using the graphical analysis method developed by Guo et al^18^. The analysis showed a strong linear correlation in *V_T_* values across all brain regions confirming that [^18^F]3F4AP and [^11^C]3MeO4AP bind to the same target (**Figure 6**). In addition, the *f_p_* values measured for each tracer and the slope estimated from the linear regression suggested that [^11^C]3MeO4AP has higher *in vivo* binding affinity than [^18^F]3F4AP and the intercept suggested higher *in vivo* specific binding. The findings of greater affinity and specific binding are likely due to the higher p*K*_a_ of [^11^C]3MeO4AP (9.1) compared [^18^F]3F4AP (7.4), which indicates that at physiological pH more of [^11^C]3MeO4AP exists in the protonated form, required to bind to the K^+^ channel^17^.

In summary, this study demonstrates that [^11^C]3MeO4AP is a promising tracer for imaging K_v_ channels in the brain, which have been closely linked to changes in demyelination and remyelination. Given the shorter half-life of [^11^C]3MeO4AP compared to [^18^F]3F4AP, this tracer has the potential to offer a viable option when multiple studies on the same subject and the same day are required.

## Materials and Methods

### Compliance

All experiments involving nonhuman primates were performed in accordance with the U.S. Department of Agriculture (USDA) Animal Welfare Act and Animal Welfare Regulations (Animal Care Blue Book), Code of Federal Regulations (CFR), Title 9, Chapter 1, Subchapter A, Part 2, Subpart C, §2.31. 2017. Experiments were approved by the Animal Care and Use Committee at the Massachusetts General Hospital.

### Radiochemistry

#### Materials

4-amino-3-hydroxypyridine and 4-amino-3-methoxypyridine were purchased from Astatech (cat# 22383 and 35474). Anhydrous solvents were purchased from Acros Organics. All other chemicals were purchased from Sigma.

#### Radiochemical synthesis of [^11^C]3MeO4AP

**Supplemental table 1** shows a summary of the conditions tested during development. *Final method*: 4-amino-3-hydroxypyridine (**1**, 2 ± 0.5 mg) was dissolved in 0.3 mL of DMSO and 4 μL of 8 N NaOH added to it. [^11^C]CH_3_I (7.4-22.2 GBq) produced in the GE Tracerlab FX MeI was bubbled into the precursor solution, sealed and allowed to react at 80 °C for 3 minutes. The reaction mixture was diluted with 5 mL of water and purified using reverse-phase HPLC (Waters XBridge C18; 5 μm, 10×250 mm; 5% EtOH in 95% sodium phosphate (10 mM, pH = 8); Flow: 3 mL/min; R_t_ ~9.3 min) or by Sep-pak purification. For Sep-pak purification the reaction mixture was diluted with 20 mL of water, loaded onto a C18 cartridge (Waters) eluted with 10% EtOH in saline. Molar activity, chemical and radiochemical purity were assessed by analytical HPLC (Waters XBridge C18; 3.5u, 4.6 x 100 mm; 5% MeCN in 95% ammonium bicarbonate (10 mM); Flow: 1 mL/min; R_t_ ~5 min). Compound identity was confirmed by coinjection with nonradioactive reference standard on HPLC.

### Imaging studies

#### Animals

Two male adult rhesus macaques (Monkey 1 and Monkey 2) were used in this study (ages: 9 and 13 years old, respectively). Animal body weights on the day of imaging were 14.5 kg on average for Monkey 1 (range 14.3 - 14.6 kg) and 16.2 kg for Monkey 2. As previously reported, one of the animals (Monkey 2) had sustained an accidental injury during a craniotomy procedure three years prior imaging^16^.

#### Animal preparation

Prior to each imaging session, animals were sedated with ketamine/xylazine (10/0.5 mg/kg IM) and were intubated for maintenance anesthesia with isoflurane (1-2% in 100% O2). A venous catheter was placed in the saphenous vein for radiotracer injection and, an arterial catheter was placed in the posterior tibial artery for blood sampling. The animal was positioned on a heating pad on the bed of the scanner for the duration of the study.

#### Magnetic Resonance Imaging

*For each monkey*, a 3-dimensional structural T1-weighted magnetization-prepared rapid gradient-echo (MEMPRAGE) was acquired using a 3T Biograph mMR (Siemens Medical Systems) for anatomical reference. Acquisitions parameters were as follows: repetition time =2,530 ms; echo time =1.69 ms; inversion time =1100 ms; flip angle = 7; voxel size = 1×1×1 mm^3^ and matrix size = 256 x 256 x 176, number of averages= 4.

#### Positron Emission Tomography Imaging

The two animals (Monkey 1 and Monkey 2) were scanned on a Discovery MI (GE Healthcare) PET/CT scanner. Each monkey underwent two baseline scans which were separated by 3 months for Monkey 1 and acquired on the same day for Monkey 2. Monkey 1 also had another scan with coinjection of unlabeled 3MeO4AP (1.16 mg/kg) which was acquired on the same day as one of the baseline scans. A CT scan was acquired prior each PET acquisition for attenuation correction. Emission PET data were acquired in 3D list mode for at least 120 min following injection of [^11^C]3MeO4AP. Radiotracer and unlabeled 3MeO4AP were administered via the lateral saphenous vein over a 3-minute infusion and followed by a 3-minute infusion of saline flush. All injections were performed using syringe pumps (Medfusion 3500). The injected dose of [^11^C]3MeO4AP at the time of injection was 292.0 ± 67.7 MBq (range 177.6 −352.8, n = 4). Molar activity (A_m_) of [^11^C]3MeO4AP at time of injection was 53.3 ± 4.1 GBq/μmol (n = 2) corresponding to injected masses of 636 ± 252 ng (n = 2). Dynamic PET data were reconstructed using a validated fully 3D time-of-flight iterative reconstruction algorithm using 3 iterations and 34 subsets while applying corrections for scatter, attenuation, deadtime, random coincident events, and scanner normalization. For all dynamic scans, list mode data were framed into dynamic series of 6×10, 8×15, 6×30, 8×60, 8×120s and remaining were 300-s frames. Final reconstructed images had voxel dimensions of 256 x 256 x 89 and voxel sizes of 1.17 x 1.17 x 2.8 mm^3^.

#### Arterial blood sampling

Arterial blood sampling was performed during the course of the dynamic PET acquisitions. Samples of 1 to 3 mL were initially drawn every 30 seconds upon radiotracer injection and decreased in frequency to every 15 minutes toward the end of the scan. [^11^C]3MeO4AP metabolism was characterized from blood samples acquired at 3, 5, 10, 15, 30, 60, 90, and up to 120 minutes. An additional blood sample of 3 mL was drawn prior tracer injection in order to measure the plasma free fraction *f_p_* of [^11^C]3MeO4AP (n = 1 for each monkey).

#### Arterial blood analysis

Radioactivity concentration (in kBq/cc) was measured in whole-blood (WB) and subsequently in plasma (PL) following the centrifugation of WB. Radiometabolite analysis was performed using an automated column switching radioHPLC system^26,27^. Eluent was collected in 1-minute intervals and assayed for radioactivity using a Wallac Wizard gamma counter (1470 model). The procedure adopted for these measurements was similar to the one described in Guehl et al^16^, except for the mobile phases used. Injected plasma samples were initially trapped on the catch column using a mobile phase consisting of either 99:1 10 mM ammonium bicarbonate pH 8 in water:MeCN or 99:1 10 mM ammonium phosphate pH 8 in water:EtOH at 1.8 mL/min and the catch column was backflushed with either 95:5 10 mM ammonium bicarbonate pH 8 in water:MeCN or 95:5 10 mM ammonium phosphate pH 8 in water:EtOH at 1 mL/min and the sample directed onto the analytical column. Standardized uptake value (SUV) radioactivity time courses in WB and in PL were generated by correcting radioactivity concentrations (C [kBq/ml]) for injected dose (ID [MBq]) and animal body weight (BW [kg]) and were calculated as SUV(t) = C(t)/(ID/BW). Measured time courses of percent parent in plasma (%PP(t)) were fitted with a single exponential decay plus a constant and individual metabolite-corrected arterial input function were derived as previously described^16^. The plasma free fraction *f_p_* of [^11^C]3MeO4AP was measured in triplicate by ultrafiltration following the procedure described in Guehl et al^16^.

#### Image registration and processing

Structural MEMPRAGE and PET images were aligned into the MRI National Institute of Mental Health Macaque Template (NMT)^21^ following the procedures extensively described in our previous work with [^18^F]3F4AP^16^. TACs were extracted from the native PET image space for the occipital cortex, parietal cortex, temporal cortex, frontal cortex, hippocampus, amygdala, striatum, thalamus, white matter and whole cerebellum. In addition, the same region of interest (ROI) as in Guehl et al^12^, manually drawn on the site of the intracranial lesion, was transposed to the native PET space for extraction of TACs.

#### Brain kinetic analysis

Extracted brain TACs were analyzed by compartmental modeling with reversible one- (1T) and two- (2T) tissue compartment models using the measured metabolite-corrected arterial plasma input function. Each model was assessed with the contribution of the WB radioactivity to the PET measurements. For quantification based on compartment models, we followed the consensus nomenclature for *in vivo* imaging of reversibly binding radioligands^28^. The primary outcome of interest was the total volume of distribution *V_T_* representing the equilibrium ratio of tracer in tissue relative to plasma and which is linearly related to the tracer binding to the target. *V_T_* was calculated as *K_1_/k_2_* for a 1T compartment model and as (*K_4_/k_2_*)×(1 + *k_3_/k_4_*) for a 2T model. The optimal compartment model was chosen on the basis of the Akaike information criterion (AIC), visual assessment of the model fits as well as robustness of the *V_T_* estimates. Graphical methods with arterial input functions, as alternative techniques for estimation of *V_T_*, were also investigated and the Logan and multilinear analysis MA1 graphical methods^29,30^ were tested. *V_T_* parametric maps were generated using the Logan method.

#### Comparison between [^11^C]3MeO4AP and [^18^F]3F4AP by graphical method

In each animal, we performed a direct comparison of the *in vivo* specific binding of [^11^C]3MeO4AP and [^18^F]3F4AP using regional *V_T_* values via linear regression, following the method described by Guo et al^18^. Briefly, in this graphical method *V_T_* of [^11^C]3MeO4AP and *V_T_* of [^18^F]3F4AP are directly related to the specific binding ratio as shown in equation 1 below,

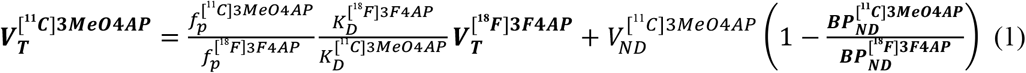

where 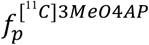 and 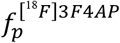 are the plasma free fractions, 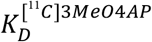 and 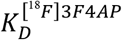 are the equilibrium dissociation constants, 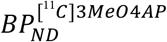 and 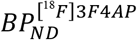 are the binding potentials (the ratio at equilibrium of specifically bound radioligand to that of nondisplaceable radioligand in tissue) for [^11^C]3MeO4AP and [^18^F]3F4AP, respectively and 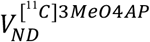 is the non-displaceable volumes of distribution of [^11^C]3MeO4AP assumed to be homogeneous across brain regions. By plotting *V_T_* of one tracer against *V_T_* of the other tracer across several brain regions with different levels of binding, this graphical method allows: (1) to determine whether the two tracers bind to the same target (assessed by the linearity or lack of linearity of the regression), and (2) to identify which compound presents the higher signal-to-noise ratio (assessed from the sign of the *y* intercept). In addition, since measurements of 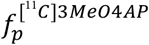 and 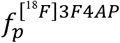 are available and both [^11^C]3MeO4AP and [^18^F]3F4AP enter the brain by passive diffusion, the slope of the linear regression provides information on the *in vivo* affinity ratio^18^. In each animal, the comparison was performed on the average *V_T_* values of the radiotracers calculated from the two baseline scans. Regional [^18^F]3F4AP *V_T_* data were obtained from Guehl et al^16^ (where Monkey 3 and 4 correspond, respectively, to Monkey 1 and 2 in the present work). Paired comparisons were plotted for the regional *V_T_* estimates in occipital cortex, parietal cortex, temporal cortex, frontal cortex, hippocampus, amygdala, striatum, thalamus, white matter and whole cerebellum.

#### Statistical analysis

All data are expressed as mean value ± one standard deviation (SD). Agreement between methods or models was assessed by computing the average measured intraclass correlation coefficient (ICC) by use of a two-way mixed-effects model with absolute agreement definition. The Akaike Information Criterion (AIC) was used to assess the relative goodness of fit between compartment models.^19^ AIC weights were computed to evaluate the probability of one model being preferred over the others.^31^ The most statistically appropriate model is the one producing the smallest AIC values and highest AIC weights. The linear correlation between *V_T_* of [^11^C]3MeO4AP and *V_T_* of [^18^F]3F4AP (equation 1) was assessed using the Pearson’s correlation coefficient *r* and the *t* distribution of the Fisher transformation was used to generate *p* values for linear regressions and intercept. A *p* value of 0.05 or less was considered statistically significant. Test-retest variability (TRT in %) of *V_T_* estimates was calculated from the two baseline scans acquired on the same day in Monkey 2 as *TRT*(% = [(*V_T,baseline2_* – *V_T,baseline1_*)/*V_T,baseline1_*]×100. The same equation was also used to calculate reproducibility of *V_T_* measurements in Monkey 1 from the baseline scans that were acquired 3 months apart.

## ACKNOWLEDGMENTS

We thank David Lee and Timothy Beaudoin at the MGH Gordon PET cyclotron facility for producing carbon-11. We thank the veterinary staff (Helen Deng and Eric McDonald) for assistance with animal handling.

## AUTHOR CONTRIBUTIONS

NJG: processed and analyzed the brain PET imaging data and blood data; RN and YZ: developed the synthesis method and synthesized [^11^C]3MeO4AP; MD and SHM: processed the blood samples and performed plasma radioHPLC analysis; GEF: contributed to the data interpretation; MDN: contributed to the study design, performed monkey brain PET scans and supervised the processing and analysis of PET data; PB: conceived the project, contributed to the study design, and supervised the entire project. NJG and PB wrote the manuscript and all authors reviewed and approved it.

