## Supplemental Information for "Radiochemical synthesis and evaluation in nonhuman primates of 3-[^11^C]methoxy-4-aminopyridine: a novel PET tracer for imaging potassium channels in the CNS"

*co-corresponding

**Author affiliations**

[1] Gordon Center for Medical Imaging, Department of Radiology, Massachusetts General Hospital and Harvard Medical School, Boston, MA

[2] Present address: Department of Radiology, University of Texas Health Science at San Antonio, San Antonio Texas, TX

*** Corresponding authors:**

Pedro Brugarolas, PhD

55 Fruit St., Bulfinch 051

Boston, MA 02114

(617) 643-4574

Marc D. Normandin, PhD

55 Fruit St., White 427

Boston, MA 02114

(617) 643-6836

### SUPPLEMENTAL TABLES

**Supplemental table 1.** Experimental conditions and yields

| **Precursor (µmol)** | **NaOH (µmol)** | **Solvent (300 uL)** | **Water (µL)** | **Temp ( ºC)** | **Time (min)** | **RCC[a] (%)** |
|---|---|---|---|---|---|---|
| 18 | 1.5 | DMF:DMSO (1:2) | 3 | RT | 3 | 13 |
| 18 | 3 | DMF:DMSO (1:2) | 3 | RT | 3 | 29 ± 2 (n = 4) |
| 18 | 25 | DMF:DMSO (1:2) | 3 | 80 | 1 | 40 ± 17 (n = 3) |
| 18 | 25 | DMSO | 3 | RT | 3 | 33 |
| 18 | 35 (KOH) | DMSO | - | 80 | 1 | 80 ± 8 (n = 3) |
| 18 | 24 | DMSO | - | 80 | 3 | 42 |
| 18 | 24 | DMSO | 4 | 80 | 3 | 35 ± 7% (n = 4) |
| 18 | 75 | DMSO | 0 | 80 | 3 | 42% (n = 1) |
| **18** | **32** | **DMSO** | **4** | **80** | **3** | **15 ± 4% (ndc RCY[b], n = 6)** |

[a] Radiochemical conversion calculated from radioHPLC chromatogram of reaction mixture
[b] Non-decay corrected radiochemical yield based on isolated product over starting [$^{11}$C]$CH_3$I activity

**SUPPLEMENTAL FIGURES**

**Supplemental Figure 1.** Radiochromatograms of plasma (PL) samples obtained at different times post injection from a representative study.

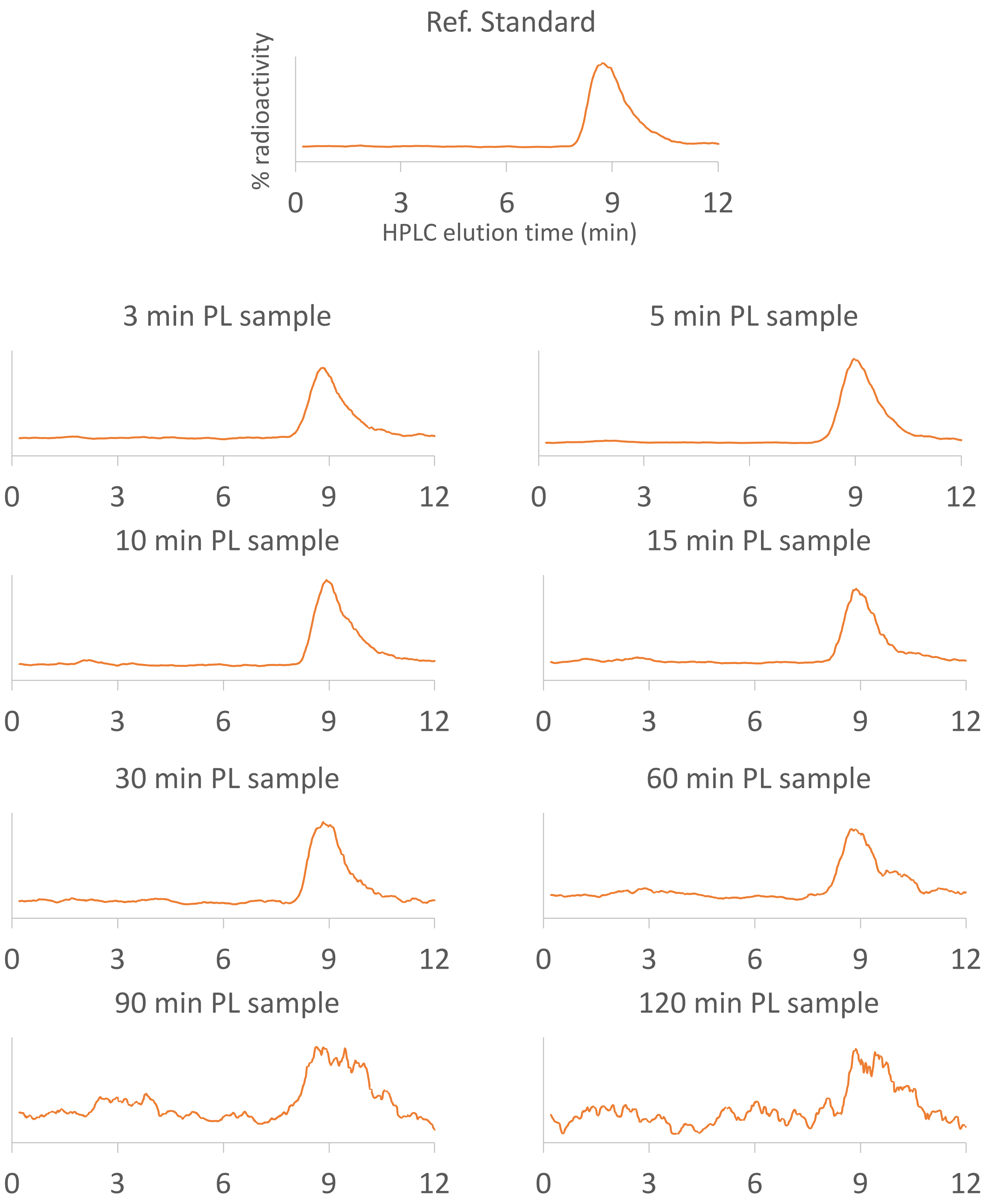
